# An integrative approach to prioritize candidate causal genes for complex traits in cattle

**DOI:** 10.1101/2024.11.11.622912

**Authors:** Mohammad Ghoreishifar, Iona M. Macleod, Amanda J. Chamberlain, Zhiqian Liu, Thomas J. Lopdell, Mathew D. Littlejohn, Ruidong Xiang, Jennie E. Pryce, Michael E. Goddard

## Abstract

Genome-wide association studies (GWAS) have identified many quantitative trait loci (QTL) associated with complex traits, predominantly in non-coding regions, posing challenges in pinpointing the causal variants and their target genes. Three types of evidence can help identify the gene through which QTL act: (1) proximity to the most significant GWAS variant, (2) correlation of gene expression with the trait, and (3) the gene’s physiological role in the trait. However, there is still uncertainty in the success of these methods in identifying the correct genes. Here we test the ability of these methods in a comparatively simple series of traits associated with the concentration of polar lipids in milk.

We conducted single-trait GWAS for ∼14 million imputed variants and 56 individual milk polar lipid (PL) phenotypes in 336 cows. A meta-analysis of multi-trait GWAS identified 10,063 significant SNPs at FDR ≤ 10% (*P* ≤ 7.15E-5). Transcriptome data from blood (∼12.5K genes, 143 cows) and mammary tissue (∼12.2K genes, 169 cows) were analysed using the genetic score omics regression (GSOR) method. This method links observed gene expression to genetically predicted phenotypes and was used to find associations between gene expression and 56 PL phenotypes. GSOR identified 2,186 genes in blood and 1,404 in mammary tissue associated with at least one PL phenotype (FDR ≤ 1%). We partitioned the genome into non-overlapping windows of 100 Kb to test for overlap between GSOR-identified genes and GWAS signals. We found a significant overlap between these two datasets, indicating GSOR significant genes were more likely to be located within 100 Kb windows that have GWAS signals compared to those without (*P* = 0.01; odds ratio = 1.47). These windows included 70 significant genes expressed in mammary tissue and 95 in blood. Compared to all expressed genes in each tissue, these genes were enriched for lipid metabolism gene ontology (GO). That is, 7 of the 70 significant mammary transcriptome genes (*P* < 0.01; odds ratio = 3.98) and 5 of the 95 significant blood genes (*P* < 0.10; odds ratio = 2.24) were involved in lipid metabolism GO. The candidate causal genes include *DGAT1*, *ACSM5*, *SERINC5*, *ABHD3*, *CYP2U1*, *PIGL*, *ARV1*, *SMPD5*, and *NPC2*, with some overlap between the two tissues.

The overlap between GWAS, GSOR, and GO analyses suggests that together these methods can identify genes mediating QTL, though their power remains limited, as reflected by modest odds ratios. Larger sample sizes would enhance the power of these analyses, but issues like linkage disequilibrium would remain.

## Introduction

Genome-wide association studies (GWASs) test the statistical association between millions of variants, such as single nucleotide polymorphisms (SNP), and a complex trait. While GWAS have been effective in identifying numerous quantitative trait loci (QTL), it is difficult to be certain which are the specific causal variants and target genes through which they influence phenotypes [1-4]. This is because of the small size of most effects, the high degree of linkage disequilibrium (LD) among nearby variants and the fact that the majority of QTL are located in non-coding regions of the genome. To identify the gene through which QTL work three types of evidence could be used: (1) genes near the most significant GWAS variant, (2) genes whose expression is correlated with the trait, and (3) genes whose physiological role is related to the trait [5]. Although none of these three types of evidence is conclusive, if they all point to the same genes that would be good evidence that the identified genes were correct. The aim of this paper is to examine the extent to which these three types of evidence agree and how often the nearest gene to QTL is the likely candidate causal gene.

Non-coding GWAS loci are likely to influence quantitative traits by modulating the expression of their target genes, in which case they are referred to as expression QTL (eQTL). There are two types of eQTL: (1) *Cis*-eQTLs are regulatory variants located near their target genes and influence expression of the gene on the same chromosome; (2) *trans*-eQTLs regulate the expression of target genes located on different chromosomes or far from the variant on the same chromosome.

As mentioned earlier, genes whose expression correlates with a trait may be genes through which QTL affect that trait. One approach to identify these genes is the transcriptome-wide association study (TWAS). TWAS combines an expression reference panel (individuals with both gene expression and genotype data) with a GWAS dataset (individuals with phenotype and genotype data) to uncover gene-trait associations [1, 2]. In fact, TWAS uses the expression reference panel to train per-gene predictive expression models, using *cis* variants i.e., SNPs typically located within 500 kb to 1 Mb of the gene’s transcription start site [TSS] [1, 2]. Predicted gene expression can then be calculated for GWAS cohorts with available genotype and phenotype data [1, 2] and the correlation between phenotype and predicted gene expression is calculated. This is commonly referred to as TWAS [1-4].

An alternative approach is to calculate the correlation between observed gene expression and genetically predicted phenotypes. This method has recently been introduced and is referred to as genetic score omics regression (GSOR) [6]. To do this, a prediction equation based on a panel of genomic variants is used to predict the trait or phenotype of interest. This linear combination of all of the SNP effects is referred to as genomic estimated breeding values (GEBVs) in animal genetics or polygenic scores (PGSs) in human genetics [7]. Usually, the largest effects on gene expression are *cis* effects, so to exploit this fact, GSOR uses the local component of this GEBV/PGS (i.e., accumulated effect of SNPs typically located within 1 Mb of the gene’s TSS) and then calculates the correlation between this local GEBV and expression of the gene. Therefore, an advantage of GSOR over TWAS is that the local GEBV/PGS is used as the response variable.

Functional knowledge of the role of the gene is the third type of evidence for the gene mediating the QTL. Functional knowledge of genes is recorded in databases such the Gene Ontology (GO) database [8, 9]. We expect that the list of genes affecting a certain trait will be enriched for functional annotations relevant to that trait. This enrichment should be clearer for traits that are physiologically simpler (e.g., human bone mineral density [5] or milk composition in cattle) than traits such as milk yield which are influenced by many physiological pathways. Here we have chosen the concentration of various polar lipids (PL) in milk as our phenotypes with the hypothesis that these traits are simpler than many other complex phenotypes.

Milk polar lipids comprising mainly phospholipids, sphingolipids and glycosphingolipids represent < 2% of total fat and are mainly located in the fat globule membrane [10]. Phospholipids phosphatidylcholine (PC), phosphatidylethanolamine (PE), phosphatidylserine (PS) and phosphatidylinositol (PI) and sphingolipids sphingomyelin (SM) are the major classes of polar lipids present in milk, whereas glycosphingolipids such as lactosylceramide (LacCer) and glucosylceramide (GluCer) are found in much lower concentration [10].

All 3 of these types of evidence are imperfect in that they are still subject to false positive or false negative findings. The extent of the overlap between the genes identified by each method provides evidence of the power of all methods, also the genes that are identified by all methods are the most likely to be correct [5].

The specific objectives of this study were: (1) to identify genomic regions associated with PL phenotypes based on a meta-analyses of multi-trait GWAS (i.e., GWAS_Meta_); (2) to identify genes from white blood cells (WBC) and mammary tissues, whose expression are significantly associated with the PL phenotypes as inferred from GSOR; (3) to perform gene list enrichment analysis of all significant GSOR genes (hereinafter referred to as GSOR hits) to identify gene ontology (GO) terms potentially involved in regulation of milk PL; and (4) to investigate the overlap between GWAS and GSOR; and (5) to investigate enrichment of GSOR hits proximal to GWAS_Meta_ signals in relevant GO terms. These sets of analyses can provide evidence of causality from three different sources of information, GWAS, gene expression, and the physiological role of those genes [5].

## Results

### Heritability estimation and GWAS for PL phenotypes

The heritability (±SE) estimated for 56 PL phenotypes using REML ranged from 0.07±0.12 to 0.7±0.13. These results are presented in Supplementary File 1. Single-Trait GWAS were performed for 56 PL phenotypes using 336 animals and 14,056,074 imputed variants (Table 1). A total of 9,923 variants demonstrated significant associations across 20 individual PL phenotypes (*P* ≤ 2.78E-5; FDR ≤ 10%), while no significant associations were identified for the remaining PL. Among the significant associations, ∼78% were located on just two chromosomes, including BTA 24 (41%), and BTA 14 (37%) (data not shown). Significant associations of GWAS_Meta_ for 56 PL phenotypes are presented in Supplementary File 2 and Manhattan plot in Figure 1. In the GWAS_Meta_, 10,063 SNPs were significant (*P* ≤ 7.15E-5; FDR ≤ 10%), of which, 13% were positioned on BTA 24, 27% on BTA 14, less than 1% on BTA 29 and 21% on BTA 26. This difference between GWAS_Meta_ and single-trait GWAS regarding the distribution of significant SNPs in various autosomes is likely due to the fact that significant associations were found for only 21 individual PL phenotypes, whereas all 56 PL phenotypes were included in the GWAS_Meta_ analyses.

**Figure 1.**
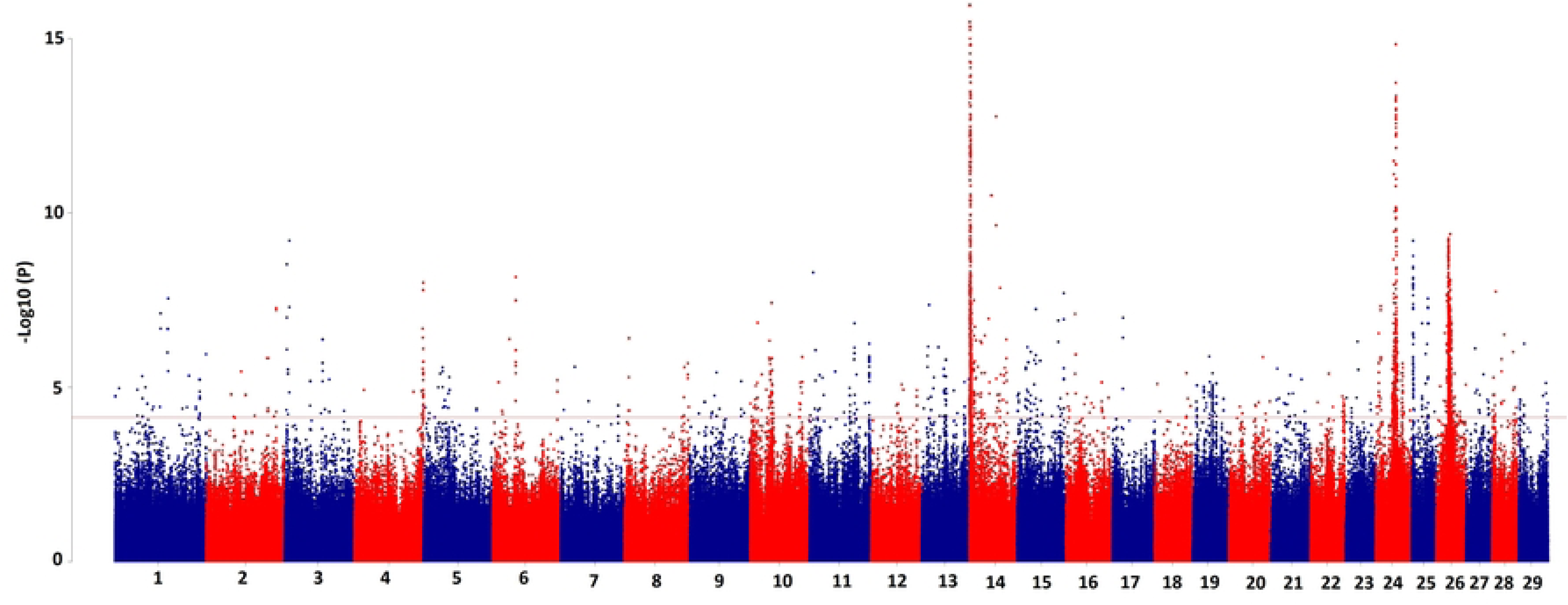
Manhattan plot for multi-trait meta-analyses GWAS for ∼14 million imputed variants and 56 species of polar lipids in milk.

**Table 1.** Data description of this study.

| Table 1. Data description of this study |  |  |  |  |  |  |
| --- | --- | --- | --- | --- | --- | --- |
| Data | Analysis | N samples Genotyped | N Variants | Phenotype | EBV | Gene Expression |
| GWAS dataset | MLM LOCO GWAS (GCTA [33]) | 336 | 14,056,074 | 56 milk PL | - | - |
| MLM LOCO GWAS results | Multi-trait GWAS [17] | 336 | 14,056,074 | 56 milk PL | - | - |
| GWAS dataset | Prediction equations (BayesR3 [48]) | 336 | 1,236,780 | 56 milk PL | - | - |
| RNA-seq WBC | Transcriptome Wide Association Study (GSOR [6]) | 143 | 1,236,780 | - | Predicted local EBV | 12,533 gene expression |
| RNA-seq Mammary | Transcriptome Wide Association Study (GSOR [6]) | 169 | 1,236,780 | - | Predicted local EBV | 12,237 gene expression |

### GSOR analysis

The correlations between the expression of 12,237 genes from mammary RNA-seq data and local GEBVs for PL phenotypes were tested to find potential associations between gene expression and individual PL phenotypes (Supplementary File 3). Collectively, we found 1,404 genes that were significantly associated with at least one PL phenotype (Supplementary File 4). The number of mammary GSOR hits across individual PL phenotypes ranged from 136 to 212, with average of 173 genes per trait (data not shown).

For the WBC dataset, the expression of a total of 12,533 genes with local GEBVs were tested (Supplementary File 5). In total, 2,186 genes were identified that were associated with at least one PL (Supplementary File 6). The number of WBC GSOR hits across individual PL traits ranged from 277 to 349 genes, with average of 314 genes per trait (data not shown).

### Functional annotation of gene lists

Using the DAVID database [8, 9], we conducted gene list enrichment analyses for 1,404 mammary GSOR hits versus 12,237 background genes, and for 2,186 WBC GSOR hits versus 12,533 background genes (Supplementary File Supplementary Files 3, 4, 5, and 6). Results are presented in Table 2. For the mammary RNA-seq data, significant enrichment was observed for the GO term lipid metabolism with 59 genes (FDR < 0.01) listed with this term and GPI-anchor biosynthesis with 8 genes (FDR < 0.05). For the WBC RNA-seq data, significant enrichment was observed for the GO terms cell adhesion (36 genes; FDR < 0.05), immune response, positive regulation of T cell-mediated cytotoxicity, and cell-cell adhesion (FDR < 0.05).

**Table 2.** Gene list enrichment analyses of mammary and white blood cell GSOR genes.

|  |  |  |  |  |
| --- | --- | --- | --- | --- |
| **Category** | **Term** | **N<sup>1</sup>** | **FDR** | **data** |
| UP_KW_BIOLOGICAL_PROCESS | Lipid metabolism | 59 | 8.4E-3 | Mammary |
| UP_KW_BIOLOGICAL_PROCESS | GPI-anchor biosynthesis | 8 | 2.8E-2 | Mammary |
| UP_KW_BIOLOGICAL_PROCESS | Cell adhesion | 36 | 3.6E-2 | WBC |
| GOTERM_BP_DIRECT | Immune response | 52 | 7.5E-3 | WBC |
| GOTERM_BP_DIRECT | Positive regulation of T cell mediated cytotoxicity | 14 | 7.5E-3 | WBC |
| GOTERM_BP_DIRECT | Cell-cell adhesion | 34 | 1.1E-2 | WBC |
| 1 Number of genes |  |  |  |  |

### Do GSOR hits agree with GWAS_Meta_ signals?

We investigated the agreement between GSOR hits and GWAS_Meta_ signals using non-overlapping windows of 100 Kb and 500 Kb. We observed a significant overlap between GSOR hits and GWAS_Meta_ signals with both tissues (*P* ≤ 0.05). These results are presented in Table 3. For example, using 100 Kb windows, we identified 24,869 non-overlapping windows, of which 839 windows included GWAS_Meta_ signals. Of our 1,404 mammary GSOR hits, 70 were found within these 839 windows with GWAS_Meta_ signals, resulting in a Fisher Exact Test *p*-value of 0.003 (odds ratio=1.47). In addition, for the WBC data with the same window size, 95 out of the total number of 2,184 GSOR hits were positioned inside the 839 GWAS-Marked windows, resulting in a *p*-value of 0.024 (odds ratio=1.28).

**Table 3.**
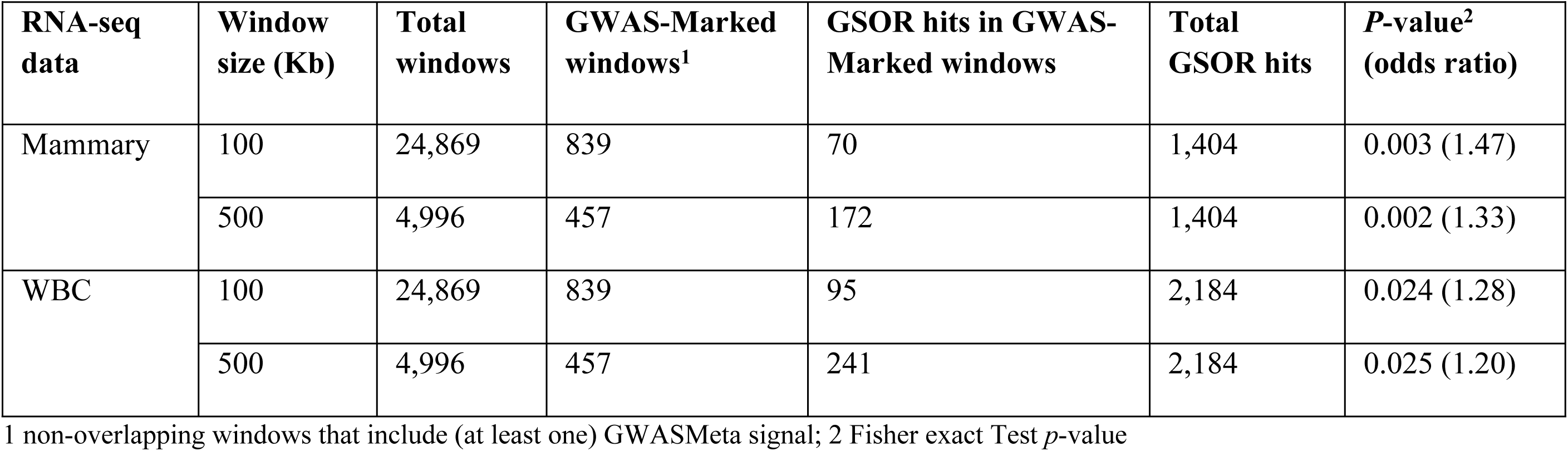
Investigation of the agreement between GSOR hits and GWAS_Meta_ signals using non-overlapping windows of various sizes.

|  |  |  |  |  |  |  |
| --- | --- | --- | --- | --- | --- | --- |
| **RNA-seq data** | **Window size (Kb)** | **Total windows** | **GWAS-Marked windows<sup>1</sup>** | **GSOR hits in GWAS-Marked windows** | **Total GSOR hits** | **P-value<sup>2</sup> (odds ratio)** |
| Mammary | 100 | 24,869 | 839 | 70 | 1,404 | 0.003 (1.47) |
|  | 500 | 4,996 | 457 | 172 | 1,404 | 0.002 (1.33) |
| WBC | 100 | 24,869 | 839 | 95 | 2,184 | 0.024 (1.28) |
|  | 500 | 4,996 | 457 | 241 | 2,184 | 0.025 (1.20) |

1 non-overlapping windows that include (at least one) GWAS<sub>Meta</sub> signal; 2 Fisher exact Test *p*-value

### Are GSOR hits located within GWAS-Marked windows enriched for the Gene Ontology term lipid metabolism?

We focused on a subset of GSOR hits located within GWAS-Marked windows and compared them with the background genes in terms of the proportion of genes involved in the lipid metabolism GO term found in each list. The results are presented in Table 4, with background genes lists in Supplementary File 3 & 5. Mammary GSOR hits located in GWAS-Marked windows were enriched with lipid metabolism GO term with all window sizes (*P* ≤ 0.01). This result for WBC data was close to significance level (*P* < 0.10) using only 100 Kb windows. The best results (odds ratio) were obtained with 100 Kb non-overlapping windows. In mammary RNA-seq data, for example, 332 of 12,237 background genes contained the lipid metabolism GO term, while seven of the 70 GSOR hits contained this GO term. This results in a *P*-value of 0.003 with odds ratio of 3.98. For the WBC data, 302 out of 12,533 background genes were listed with the lipid metabolism GO term, while five out of 95 GSOR hits within GWAS-Marked windows contained the same GO term (*P* = 0.08; odds ration=2.24).

**Table 4.** Enrichment of those GSOR hits that are located within GWAS-Marked windows in Lipid metabolism GO compared to background genes, using different non-overlapping window sizes.

| RNA-seq data | Window size (Kb) | GSOR hits within GWAS-Marked windows<sup>1</sup> |  | Background genes |  | *P*-value<sup>3</sup> (odds ratio) |
| --- | --- | --- | --- | --- | --- | --- |
|  |  | Involved in<sup>2</sup> | Not involved in<sup>2</sup> | Involved in<sup>2</sup> | Not involved in<sup>2</sup> |  |
| Mammary | 100 | 7 | 63 | 332 | 11,905 | 0.003 (3.98) |
|  | 500 | 11 | 161 | 332 | 11,905 | 0.008 (2.44) |
| WBC | 100 | 5 | 90 | 302 | 12,231 | 0.08 (2.24) |
|  | 500 | 9 | 232 | 302 | 12,231 | 0.19 (1.57) |

<sup>1</sup> GWAS-Marked windows are non-overlapping windows of different length that contain at least one GWASMeta signal; <sup>2</sup> Number of genes involved in and not involved in Lipid metabolism GO, respectively; <sup>3</sup> Fisher Exact Test *p*-value.

Therefore, GSOR hits from mammary and WBC data that were located in GWAS-Marked windows and are known to be involved in lipid metabolism (based on GO annotation) were potential candidate causal genes as they were supported by three sources of information (Table 5). These candidate causal genes include *DGAT1, SERINC5, PIGL, CYP2U1, ABHD3, CSM5, and ARV1* from mammary gland and *DGAT1, PIGL, CYP2U1, SMPD5* and *NPC2* from WBC data. Figure 2 illustrates two examples (*DGAT1* and *SERINC5*), showing the mammary GSOR hits and background genes in their proximity, along with the Manhattan plot from GWAS_Meta_.

**Figure 2.**
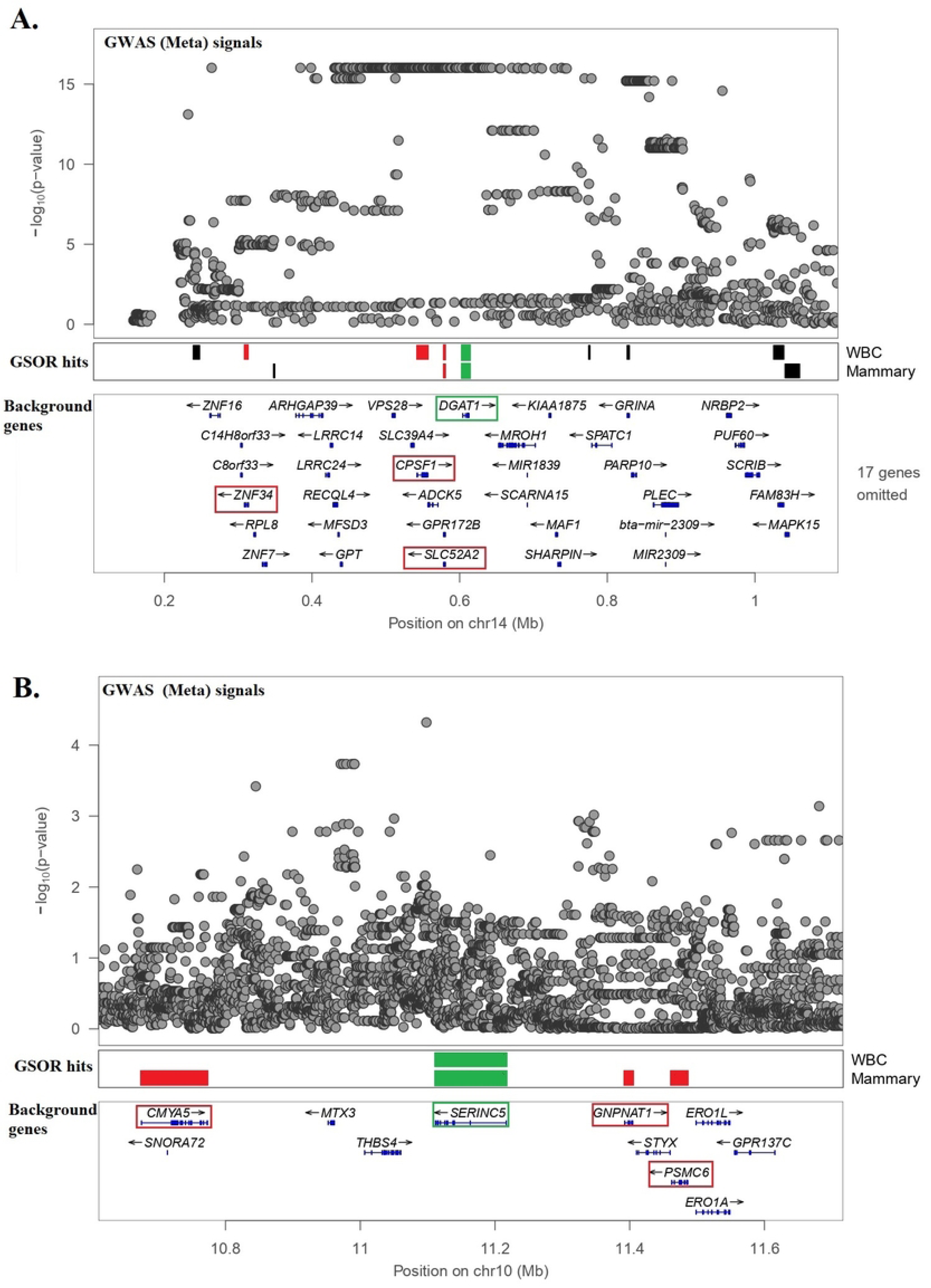
GWAS and GSOR signals surrounding two of the prioritized genes, *DGAT1* and *SERINC5*. The first level of the plot shows the Manhattan plot for GWAS meta-analysis, the second level shows the genes whose expression in mammary gland and/or WBC data were significantly associated with at least one PL phenotype (genes highlighted with green are potential candidate causal genes), and the third level plot represents background genes expressed in mammary data.

**Table 5.**
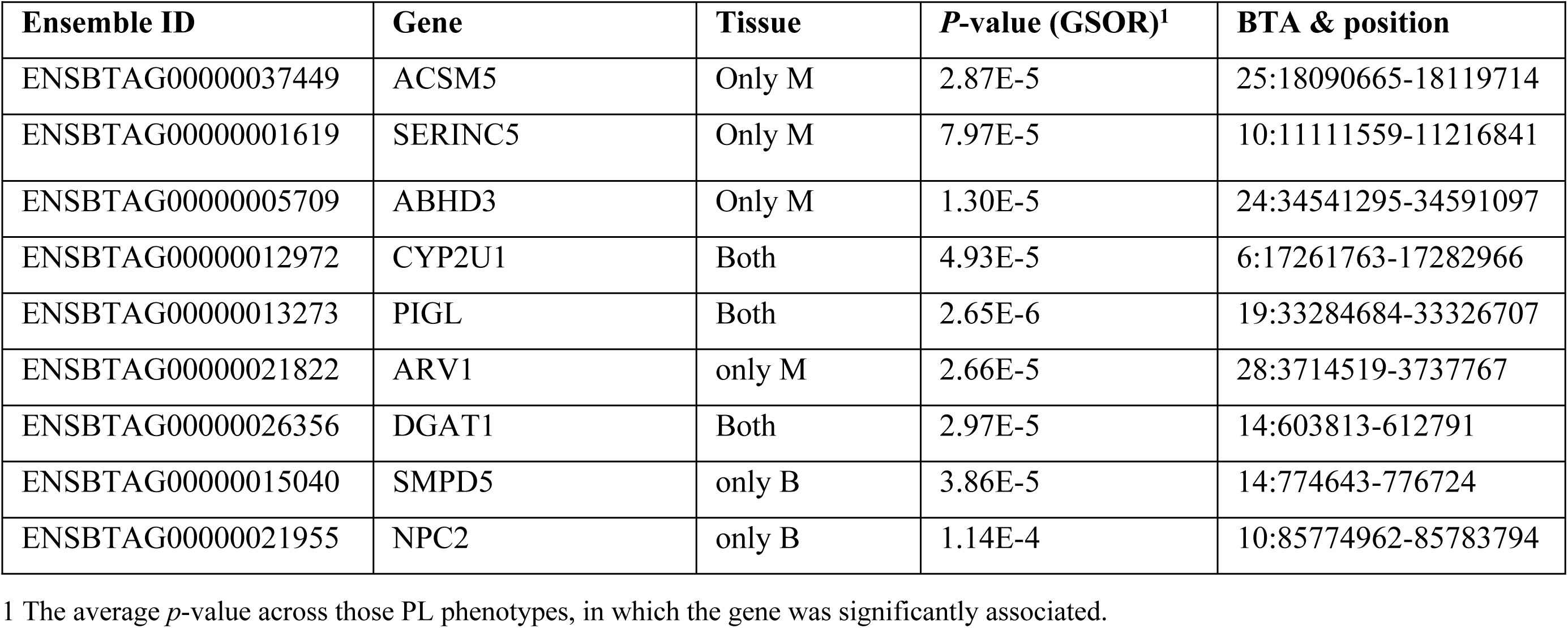
Candidate causal genes for PL traits that are a subset of GSOR genes located in GWAS-Marked non-overlapping windows of 100 Kb and involved in Lipid metabolism GO term.

## Discussion

GWAS have identified many variants associated with complex traits e.g., in human height [11] or in livestock production traits [12], etc. If the causal variant is located in a coding region, it may directly point to the associated gene. However, studies have reported that most variants associated with complex traits by GWAS are located in non-coding regions [13, 14] and therefore have unknown functions. QTL in a non-coding regions can affect phenotype by regulating gene expression (i.e., eQTL) [15, 16]. The challenge, largely due to LD, is not only to pinpoint the actual causal variants in non-coding regions, but also to find the target genes through which they affect phenotype [2, 4, 6]. Therefore, TWAS and other methods have been developed to prioritize causal genes at GWAS loci [1].

In this study, we integrated GWAS_Meta_ with GSOR analyses to identify potential candidate causal genes affecting complex trait phenotypes (i.e., concentration of PLs in milk). Using meta-analyses on 56 single-trait GWAS (GWAS_Meta_), 10,063 associations were identified. The GWAS_Meta_ approach was used because meta-analyses can enhance the power to detect genetic variants by leveraging the shared genetic architecture across correlated traits [17]. Next, GSOR was performed to identify significant associations between gene expression and local GEBVs for PL phenotypes. Our GSOR analysis revealed 2,186 and 1,404 genes from WBC and mammary transcriptome were significantly associated with at least one PL phenotype. However, not all of these gene-trait associations are causal. TWAS methods (including GSOR) often detect multiple significant genes per locus [2, 5].

Several factors contribute to TWAS cause false positives [2]. For instance, when an eQTL affecting phenotype is in LD with another eQTL, this will cause a correlation between expression of both genes and the trait, but the relationship is not causal for the second gene [2]. In our study, both *DGAT1* and *SLC52A2* genes were GSOR hits, with the former being a very well-known causal gene affecting milk fat [18-22] (Figure 2a). It seems that *SLC52A2* is probably a false GSOR hit because its expression is highly correlated (r=0.5, *p*=2.8E-12) with expression of *DGAT1* in mammary gland. Another example is the correlation of expression between *SERINC5* and *PSMC6* gene (r= -0.34; *p* = 3.7E-6); while the former was one of our prioritized candidate causal genes, the latter would be more likely a false hit (Figure 2b). Thus, assuming that gene expression mediates the genetic impact on complex traits, GSOR or TWAS associations do not offer direct evidence of causal links between gene expression and these traits. Instead, they represent associations between expression levels and phenotypes [2]. Therefore, additional sources of evidence are required to prioritize candidate causal gene-trait associations.

We assessed the agreement between GSOR hits and GWAS_Meta_ signals. By partitioning the genome into non-overlapping windows of different sizes including 0.1 and 0.5 Mb, we demonstrated that GSOR hits were significantly more likely to be located within GWAS-Marked windows. This significant overlap supports the hypothesis that GSOR hits include causal genes which mediate the effect of GWAS_Meta_ signals on the PL phenotypes.

If GSOR hits tagged by GWAS_Meta_ signals (i.e., those genes that are located within GWAS-Marked windows) include causal genes, they should show enrichment for biologically relevant GO. Milk PLs are promising target phenotypes to test this hypothesis. Traits like milk yield are not simple, and many pathways likely influence the final phenotypes. However, identifying enriched pathways should be easier with physiologically simple phenotypes like milk PLs. Additionally, the medium to high heritability of most PL phenotypes increases the likelihood of detecting genetic associations, even with smaller sample sizes. We tested this hypothesis by investigating enrichment of lipid metabolism GO terms for genes that were GSOR hits located within GWAS-Marked windows compared to total background genes. Results indicated a significant enrichment of these GSOR hits for lipid metabolism GO (Table 4), particularly in mammary tissue, where the 100 Kb windows showed the strongest associations (odds ratio=3.98; *P* = 0.003). Seven of the 70 mammary GSOR hits located within GWAS-Marked windows contained the lipid metabolism GO term (∼10%). These genes are potential candidate causal genes for PL phenotypes.

The mammary GSOR hits that were prioritized as candidate causal genes were *DGAT1*, *ABHD3*, *SERINC5*, *CYP2U1*, *PIGL*, *ARV1*, and *ACSM5*. Our findings revealed that the expression of *DGAT1* in mammary gland is associated with a different PL phenotype including phosphatidylserine, sphingolipids sphingomyelin, phospholipids phosphatidylcholine, phosphatidylinositol and glucosylceramide phenotypes. The *DGAT1* gene has been reported to account for 30-40% of the phenotypic variance of milk yield and composition in cattle [23, 24]. Although a protein-coding mutation for this gene has already been identified [24, 25], the present study and previous ones [20, 23, 26] revealed *cis*-regulatory effects for *DGAT1*, possibly attributable to multiple causal mutations. GSOR analysis of mammary tissue found *ABHD3* on BTA24 to be significant, we observed the 2^nd^ most significant GWAS_Meta_ peak surrounding this gene. In cattle, a low-resolution GWAS found a broad region on BTA 24 including *ABHD3*, among many other genes, associated with concentration of milk fatty acids [27]. In human, this gene was associated with circulating phospho- and sphingolipid concentrations [28], with a recent GWAS on plasma lipidome reporting a missense mutation causing this association [29]. In a metabolomics study on *ABHD3*, this gene was annotated as a lipase that specifically targets medium-chain and oxidatively truncated phospholipids, establishing its physiological role in lipid metabolism [30]. This is the first time that a *cis*-regulatory mechanism linking expression of the *ABHD3* gene in mammary gland to the concentration of PLs in milk has been reported. Our findings showed that the expression of *ABHD3* gene in mammary gland is associated mainly with sphingolipids sphingomyelin and phospholipids phosphatidylcholine, but also associated with phosphatidylethanolamine and phosphatidylserine. Another gene is *ACSM5* (Acyl-CoA Synthetase Medium Chain Family Member 5) on BTA 25, whose expression in mammary gland is associated mainly with SM phenotypes, but also with one of the phosphatidylethanolamine phenotypes. *ACSM5* catalyses the activation of fatty acids by CoA to produce an acyl-CoA, which is the first step in fatty acid metabolism. *ACSM5* is involved in the fatty acid biosynthetic process and the acyl-CoA metabolic process (https://www.genecards.org/). The *ARV1* gene on BTA28, whose expression in mammary gland is associated with glucosylceramide phenotype, is listed with several GO terms including sphingolipid, cholesterol, bile acid metabolism and cholesterol and sterol transport. It has been reported that Yeast cells lacking the *ARV1* gene harbor defects in sphingolipid metabolism [31].

Our study has some limitations. First, despite the relatively simple physiology and higher heritability of PL traits, the sample size used to predict GEBVs and estimate their correlations with gene expression was small. This could have limited the power of BayesR3 to identify variants with smaller effect on PL concentration. We recommend testing the methods presented here with a larger dataset. Furthermore, a significant portion of the heritability of complex traits can be linked to *trans*-eQTLs, eQTL located on different chromosomes or more than 5 Mb away [32], and these were not included in this study. However, study on *trans*-eQTLs requires a large number of expression reference samples to ensure adequate statistical power.

In conclusion, the significant overlap between the genes identified by GWAS, GSOR and GO indicated that all three methods have some power to identify genes mediating QTL. However, the odds ratios in Tables 3 and 4 are not very high, so the power of these methods is limited. Power could be increased by larger sample size, but we anticipate that some problems, such as LD, would persist. However, the combination of methods does give a list of candidate genes with fewer false positives.

## Material and Methods

### GWAS data description

Phenotypic data for 336 Australian Holstein cows including records for concentration of 59 species of PLs in milk were available. Polar lipids were extracted from raw milk as previously described [10]. Internal standard (PS 34:0) was added prior to lipid extraction. An Agilent 1290 UPLC system coupled to an LTQ-Orbitrap MS (Thermo Scientific) was used for polar lipid quantification. Chromatographic separation of polar lipids was achieved using a Luna HILIC column (250×4.6 mm, 5 µm, Phenomenex) maintained at 30 °C. The mobile phase was composed of 5 mM aqueous ammonium formate (A) and acetonitrile containing 0.1% formic acid (B). The flow rate was 0.6 mL/min with a gradient elution of 2 to 21% A over 25 min. The injection volume was 5 µL. The detection of lipids was by LTQ-Orbitrap mass spectrometer (Thermo Scientific) operated in electrospray ionization positive (for most polar lipid classes) or negative (for analysis of PI) Fourier transform mode. The resolution was set to 60,000 for both positive and negative modes. Identification of lipid species present in milk was performed as previously reported [10]. Quantification of selected polar lipid species was based on peak area of parent ions after normalization by the internal standard.

Heritability (h^2^) was estimated for these traits using --reml command in GCTA [33]. Three PLs were excluded due to zero heritability, leaving 56 PLs for GWAS analysis (Table 1). Records with ±4 SD from the mean phenotype were excluded from the analysis.

Of these 336 cows, 181 were genotyped with Standard 50K SNP chips, 17 cow genotyped with High Density (HD) 700K, the remainder with Low Density ∼7.5K. Genotype data for the 336 cows was imputed to the whole genome sequences (WGS) level (Table 1). Minimac3 [34] was used to impute genotypes with Run7 of the 1000 Bull Genomes project as reference population [35]. The details of the imputation are described in [36].

### Expression reference panel data

Two different sets of transcriptomics data were used from WBC and mammary tissues.

WBC gene expression was obtained from 143 Australian Holstein animals, a subset of a larger multibreed dataset. The processing of samples, RNA extraction, library preparation, RNA sequencing, etc. were described in detail in [32, 37, 38] (Table 1). This data did not overlap with the GWAS dataset. Imputation to WGS genotypes for these animals was conducted as described above for the GWAS animals.

Mammary gene expression data was obtained for 169 New Zealand Holstein cows [39-41](Table 1). For this dataset, 12,622,468 variants were imputed using 1,298 reference animals (including 306 Holstein-Friesian, 219 Jersey, 717 HF×J, 56 other breeds) as described in [42].

### GWAS analysis

Variants with low imputation accuracy (Minimac R^2^ < 0.5), minor allele frequency (MAF) < 0.01, and variants deviating from Hardy-Weinberg Equilibrium with χ^2^ *p*-value < 1E-6 were excluded. Single-trait GWAS were conducted for 56 PL phenotypes (Supplementary File 1) using 14,056,074 imputed variants. For GWAS we used a mixed linear model leaving-one-chromosome-out (MLM-LOCO) approach implemented in GCTA software [33, 43]. The model used is as follows:

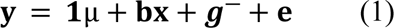

where **y** is a vector of records for phenotypic values; **1** is a vector of ones; µ is the mean of the trait; **b** is the additive allelic substitution effect of the SNP being tested; **x** is a vector of allele dosages (coded as 0, 1 or 2); ***g***^-^ is a vector of polygenic effects with 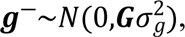 where **G** is the genomic relationship matrix [44] calculated using SNP 50K genotypes, excluding variants located on the chromosome harbouring the test SNP (LOCO approach); 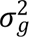 is the additive genetic variance explained by the 50K SNPs; and ***e*** is the vector of residuals with 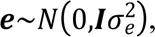 where **I** is an identity matrix and 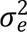 is residual variance.

### Multi-trait GWAS meta-analysis

We performed a meta-analysis using outputs from 56 single-trait GWAS (hereinafter referred to as GWAS_Meta_) [17]. The following formula for GWAS_Meta_ was used [17]:

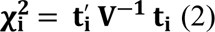

where **t**_**i**_is a n×1 vector of signed t-values of *SNP_*i*_*, and n is the number of traits used; **t**^′^_**i**_ is transpose of **t**_**i**_ and **V**^**-1**^ is the inverse of a *n* × *n* correlation matrix where the correlation between two traits is the correlation over the 14,056,074 estimated SNP effects (signed t-values) of the two traits; 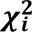 is the chi-squared statistics with *n* degrees of freedom for the i^th^ SNP; the *p*-value for the i^th^ SNP was calculated using *pchisq* R function [45] with *n* degrees of freedom [17]. To account for multiple testing, obtained *p*-values were adjusted using Benjamini-Hochberg method [46] implemented in *p.adjust* R function [45]. Variants with FDR ≤ 0.10 were treated as significant in GWAS_Meta_.

### Predicting GEBVs for expression reference panel cows

We performed LD-pruning on genotypes of GWAS cows using PLINK v1.9 [47] with parameters --indep-pairwise 5000 500 0.95 to exclude variants that were in strong LD (r^2^ > 0.95). The LD-pruned GWAS dataset with 1,236,780 SNP was used to train models with BayesR3 software [48] to predict GEBVs for the PL traits on the expression reference panel cows (Table 1). Individuals from the expression reference panel had the same set of 1,236,780 SNP genotypes. For each individual PL, the following model was used:

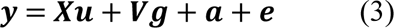

where ***y*** is an *n* × 1 column vector of phenotypic records, in which *n* is number of records; ***X*** is an *n* × *m* incidence matrix, **u** is *m* × 1 vector of fixed effect and *m* is the number of fixed effects including the combined effect of batch and year (nine levels); **V** is the coded genotype, representing the observed genotypes of each individual; **g** is a vector of SNP effects; **a** is vector of random genetic effects not explained by the SNPs with polygenic variance represented as 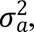 in which 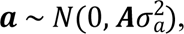 and **A** is the relationship matrix; and **e** is the residual term. BayesR3 was run with 50,000 MCMC iterations and 25,000 burn in. In the BayesR3 model, the SNP effects follow a mixture of four normal distributions with zero mean and additive genetic variances of zero, 0.0001, 0.001, and 0.01 times the genetic variance. Starting values for proportions of the four SNP effect distributions were defined as 0.994, 0.0055, 0.00049, and 0.00001 respectively.

Once SNP effects on an individual PL phenotype were estimated, local GEBVs for cows in the expression reference panel, corresponding to a specific gene, were calculated using the effect of SNPs positioned within ±1 Mb of that gene’s TSS.

### Genetic Score Omics Regression (GSOR)

For an individual PL and RNA-seq dataset (i.e., tissue), the following per-gene GSOR model was applied:

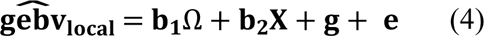

where 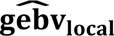 is a *m* × 1 vector of local GEBVs (corresponding to a gene); Ω is a *m* × 1 vector of that gene’s expression; and m is the number of animals; **b**_**1**_ is the regression coefficient of the 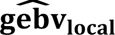 on Ω; **X** represents a design matrix for fixed effects, and **b**_**2**_ is the vector of fixed effect (for the RNA-seq data); and **g** is a vector of random polygenic effects (for the RNA-seq data) with 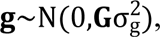 where **G** is the genomic relationship matrix [44] calculated using 50K SNP genotypes, and 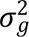 is the additive genetic variance explained by the 50K SNPs; ***e*** is the vector of residuals with 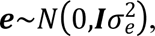 where **I** is an identity matrix and 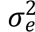 is residual variance.

Models for the WBC RNA-seq dataset were fitted using the experiment (with 6 levels) and days in milk (DIM) as categorical and quantitative fixed effects, respectively. Fixed effects were not available for the mammary RNA-seq dataset. Once associations between all the genes within an RNA-seq dataset and an individual PL (i.e., local GEBV for cows corresponding to that gene) were estimated, *p*-values were adjusted for multiple testing using Benjamini-Hochberg method [46]. Genes with FDR ≤ 0.01 were considered significant.

### Functional annotation analyses

Gene-set enrichment analyses were performed for each tissue separately using GSOR genes that were significant across 56 PL phenotypes, where the total number of genes (tested genes) within each RNA-seq dataset was used as background genes. To do this, DAVID (The Database for annotation, Visualization, and Integrated Discovery) bioinformatic tool [8, 9] was used and biological terms with FDR ≤ 0.05 were regarded as significant.

### Do GSOR hits agree with GWAS_Meta_ signals?

To test this hypothesis, the genome was partitioned into 100 Kb and 500 Kb non-overlapping windows, and windows containing at least one significant GWAS_Meta_ SNP were identified (hereinafter referred to as GWAS-Marked windows). We then counted the GWAS-Marked windows and GSOR hits found within these windows. A GSOR hit is considered to be within a GWAS-Marked window if that gene’s start position falls within that window. These values were compared to the total number of non-overlapping windows and the total number of GSOR hits. Fisher’s Exact Test was used and a *P*-value ≤ 0.05 was considered significant.

### Are GSOR hits found within GWAS-Marked windows enriched for relevant GO terms?

To test this hypothesis, the proportion of lipid genes (i.e., genes involved in Lipid metabolism GO) were compared in two gene lists: (1) GSOR hits found within GWAS-Marked windows (described above) versus (2) the total number of tested genes in that RNA-seq data (i.e., background genes). Each gene list was categorized into two groups including genes “involved in” and genes “not involved in” lipid metabolism GO. Fisher’s Exact Test was used to investigate the significance of difference between proportions in these categories for the two gene lists, and a *P*-value ≤ 0.05 regarded as significant.

## Ethics approval and consent to participate

No new animal experiments were undertaken for this study, all data was taken from previously published studies. Refer to materials and methods for details.

## Data Availability

All gene expression data was taken from previously published studies as detailed in the materials and methods. The Polar Lipids data used in the study though taken from a previously published study (see materials and methods) had not been publicly released. It will be publicly available upon acceptance of this paper.

## Acknowledgments

We thank Dr. Bolormaa Sunduimijid for imputation of sequence variants for cattle with Polar Lipid phenotypes and the partners from Run7 of the 1000 Bull Genomes Project for data access.

## Supplementary Files

Supplementary File 1. REML based heritability estimations for 56 milk polar lipid phenotypes

Supplementary File 2. Significant SNPs for meta-analysis of genome-wide association study (GWAS) for 56 single-trait GWAS

Supplementary File 3. Ensemble ID for background mammary genes

Supplementary File 4. The 1,404 GSOR hits identified for mammary transcriptome

Supplementary File 5. Ensemble ID for background white blood cells genes

Supplementary File 6. The 2,186 GSOR hits identified for white blood cells transcriptome

Supplementary File 7. Genes involved in Lipid metabolism GO identified using background mammary genes

Supplementary File 8. Genes involved in Lipid metabolism GO identified using background white blood cells genes

